# DepoCat: Interactive database of experimentally verified phage depolymerases

**DOI:** 10.64898/2026.09.11.750917

**Authors:** Sebastian Olejniczak, Aleksandra Otwinowska, Maria Pozniak, Zuzanna Drulis-Kawa

## Abstract

*Klebsiella* phage depolymerases degrade polysaccharide capsules and exhibit narrow substrate specificity for particular capsular types. Despite a growing number of experimentally characterized enzymes, these data remain scattered throughout the scientific literature, while existing protein sequence repositories are dominated by entries with computationally assigned, unverified functional annotations. Here we present DepoCat, the first interactive database of phage depolymerases with experimentally verified function and specificity, available at http://depocat.uwr.edu.pl. The database currently contains 131 proteins meeting rigorous inclusion criteria, spanning 75 distinct capsular types. Each entry integrates experimental and computational resources. The web interface provides an integrated Classifier tool with two search modes: sequence-based search and structure-based search – enabling preliminary structural classification and inference of putative substrate specificity of newly identified depolymerases. We demonstrated the utility of both modes on a set of 17 experimentally verified non-Klebsiella phage depolymerases, for which structural analysis enabled unambiguous class assignment in almost all cases despite low or undetectable sequence similarity to the database reference dataset. DepoCat constitutes a publicly accessible resource supporting research into the structural diversity and sequence-structure-specificity relationships of phage depolymerases, while also facilitating the identification of candidates for therapeutic and diagnostic applications.

## Introduction

Bacterial exopolysaccharides (capsules or lipopolysaccharides (LPS)) are a key virulence factor for many Gram-negative bacteria, protecting cells from recognition by the innate immune response, adaptive immune recognition, and antimicrobial agents (1,2). In *Klebsiella pneumoniae*, 163 distinct capsular loci have been identified to date; however, the monosaccharide composition and architecture have been defined for at least 79 of them, with the rest remaining unknown (3–7). The high diversity of capsular types, together with their direct link to pathogenicity, poses a significant challenge for the development of effective anti-virulence therapeutic strategies.

As natural bacterial predators, bacteriophages have evolved specific receptor-binding proteins incorporating depolymerases that enable enzymatic degradation of bacterial exopolysaccharides as a first step of infection (8,9). Phage depolymerases exhibit narrow substrate specificity, making them precise therapeutic agents (10–12) that sensitize the bacterium to the host immune system and treatment (9,13). The depolymerase application has been demonstrated in mouse models of *K. pneumoniae* infections, where treated animals showed higher survival rates compared to untreated controls (14–19).

Beyond their therapeutic potential, phage depolymerases might find application in microbiological diagnostics, serving as tools for rapid typing based on substrate specificity and offering an alternative to classical antibody-based serological methods. Notably, in certain cases their substrate specificity may surpass that of antibodies, allowing highly accurate identification of distinct capsular polysaccharide types (17,20). It was also shown that phage depolymerases can serve as promising prototypes for next-generation anti-virulence and vaccine strategies against multidrug-resistant *K. pneumoniae*, since oligosaccharide products generated during degradation can modulate immune responses (21).

Exploring phage-derived enzymes is itself important in understanding phage biology. Since enzymatic degradation of the capsule constitutes a key step in phage adsorption to the bacterial cell, depolymerase specificity determines the host range of a given phage and provides insight into the molecular basis of phage-host interaction at the receptor-ligand level (22,23). From an evolutionary perspective, depolymerases are an exceptional model for studying phage-bacteria co-evolution. The high diversity of bacterial capsules exerts selective pressure that drives phages to evolve new enzyme repertoires capable of recognising and degrading new capsule variants. This dynamic offers insight into the mechanisms underlying phage adaptation to changing hosts (10,24).

Although the number of experimentally characterized phage depolymerases is increasing, the relationships between amino acid sequence, structure, and substrate specificity remain poorly understood. The available pool of confirmed data is still too limited for systematic comparative analysis (11), and information regarding their experimental characterisation is scattered across individual publications, lacking a central, searchable, and comparable resource. While large protein sequence repositories such as GenBank and UniProt contain huge numbers of sequences and predictions, the vast majority have not been experimentally tested for enzymatic activity and substrate specificity to understand the sequence-structure-specificity relationship.

The rate of incorrect molecular function annotation in GenBank NR and TrEMBL averages 5-63% depending on the enzyme superfamily, with misannotation exceeding 80% in at least one database for 10 of the 37 families examined (25). This problem has not been diminished in subsequent decades. Rembeza and Engqvist experimentally verified that at least 78% of sequences in a studied enzyme class are misannotated, and a broader computational analysis of the BRENDA database revealed that nearly 18% of all enzyme sequences share no detectable similarity to experimentally characterised representatives of their assigned class, including enzymes of established industrial relevance (26). The issue is further compounded by the interconnected nature of public repositories, where erroneous annotations propagate across linked databases as new sequences automatically inherit functional assignments from existing records (27). Although the problem was recognised over a decade ago, comprehensive analyses of its scale in the era of large-scale genome sequencing remain scarce, and available experimental evidence indicates that it has not abated (26,28). The consequences of propagating unverified functional annotations extend well beyond the misidentification of therapeutic candidates. In evolutionary and comparative genomics, misannotated sequences can distort the perceived diversity and phylogenetic distribution of enzyme families, generating erroneous conclusions. In biotechnology and synthetic biology, engineering efforts aimed at modifying substrate specificity depend on accurate reference sequences. Building upon unvalidated entries risks propagating errors into engineered variants. Finally, in epidemiological and diagnostic contexts – such as depolymerase-based typing panels – experimentally confirmed substrate specificities are a prerequisite for the reliable assignment of capsular types to clinical isolates (29).

The absence of a dedicated, curated database consolidating experimentally validated phage depolymerases into a single, publicly accessible resource represents a significant gap in the bioinformatics repositories available to researchers working with phage biology, phage-bacteria interactions, virus evolution, biotechnology, therapy, and diagnostics.

Here, we present DepoCat, the first interactive database of Klebsiella phage depolymerases (http://depocat.uwr.edu.pl) as a starting point for a curated depolymerase database and tool. Each entry in the database features experimentally verified enzymatic activity and defined substrate specificity. The database integrates experimental and computational data, including structural models, domain annotations, and a novel structural classification system (12). The web interface also provides a Classifier tool with sequence and structure search modes, which enable preliminary classification and inference of putative substrate specificity for newly identified phage depolymerases.

## Materials and Methods

### Database compilation

The DepoCat database was compiled using 129 experimentally characterised Klebsiella phage depolymerases described in our recent publication (12) and two additional enzymes described separately (30). The full list of proteins included in the DepoCat database is incorporated in Supplementary Table S1. 90 out of the 131 enzymes were screened against a big collection of *Klebsiella spp*. strains belonging to K1-K82 defined serotypes or genetically classified to KL101-KL186 with some exceptions (KL106, KL129, KL147, KL150, KL152, KL154, KL156, KL157, KL159, KL160, KL162, KL171, KL172, KL175, KL176, KL179, KL180, KL182, KL185) (12).

The database consists of proteins meeting the following criteria: (I) confirmed recombinant production, (II) experimentally validated enzymatic activity, and (III) experimentally determined substrate specificity to one or more defined *Klebsiella* capsular types. We exclude proteins identified solely through computational prediction or sequence homology without functional confirmation.

Each database entry contains two types of data (experimental and computational). Experimental data include confirmation of recombinant production with enzymatic activity, substrate specificity, the full panel of tested capsular types, and the phage’s name, morphotype, life cycle, and bacterial host. Computational data comprise nucleotide and amino acid sequences, molecular weight, and structural models generated as homotrimers using AlphaFold3 (31,32), presented in two variants, one coloured by domain organization and the other by pLDDT prediction confidence score. Additional calculated data include average prediction quality metrics: pLDDT, pTM, ipTM, and PAE. The domain organisation was determined based on the structural models curated manually using PyMOL Open-Source (v3.1.0)(33). Phages, a source of proteins, were classified according to current ICTV taxonomy (34)(MSL#41, 20 March 2026). Each depolymerase is also assigned to a structural class and subclass based on a novel classification system (12), comprising five structural classes and 14 subclasses through manual analysis of domain architecture combined with pairwise structural comparisons using the US-align (v20260328) software (35).

### Database design and web application implementation

All protein data in the DepoCat database are stored in a structured JSON (JavaScript Object Notation) file, which serves as the central data source for the entire application. Structural models are stored in GLB format for the Google 3D model-viewer and in PDB format for the structure search. The application frontend was built using React (v19.2.6) and Vite (v8.0.12), with client-side navigation handled by React Router (v7.15.1). Visualisation of 3D structural models is implemented using the Google 3D model-viewer web component. The backend was implemented using the FastAPI framework (Python 3.10.16), which handles serving static frontend files and the REST API. Long-running bioinformatics computations are performed asynchronously using the Celery (v5.6.3) task queue system and results backend based on Redis (v7.0.15). The application is deployed on a university server running Linux, with uvicorn as the ASGI server.

### Classifier tool

DepoCat features an integrated Classifier tool offering two search modes: sequence search and structure search. The sequence search employs the BLASTP program (v2.17.0+)(36) with the following parameters: an E-value threshold of 0.05, a maximum of 131 target sequences, JSON output format, and the BLOSUM62 scoring matrix. The structure search utilises the US-align (v20260328) software (35). The user-submitted structural model is compared against each of the 131 reference structural models stored in the database (v2.0; 26 June 2026). The resulting structural superposition file is combined with the reference structural model into a single PDB file containing two MODEL records: MODEL 1, corresponding to the transformed query structural model, and MODEL 2, corresponding to the reference structural model. The 3Dmol.js library Visualisation the structural superpositions (37).

### Sequence and structural comparisons of depolymerases

BLASTP (v2.17.0+)(36) with default settings was used for sequence comparisons. Protein structural models were obtained from the PDB database (RCSB.org) (38) and compared using USalign (v20260328) (35). PyMOL Open-Source (v3.1.0) was utilized for structural alignments and visualisations (33).

## Results

### Web interface

The DepoCat database is available at http://depocat.uwr.edu.pl as a free web application that requires neither registration nor login. The interface provides two entry points for browsing the database: by K/KL type specificity and by structural class. The detail page for each protein presents a comprehensive dataset arranged into thematic panels. The domain organisation panel displays a segmented diagram proportional to the length of each domain within the amino acid sequence, accompanied by a table listing domain names, positions, and lengths. The tested capsule type panel displays all known K/KL types using colour-coded markers: green for types against which the protein shows activity, red for types tested but showing no activity, and grey for untested types. Two interactive structural model panels allow visualisation of models coloured according to domain organisation and pLDDT prediction confidence score. Structural models for each depolymerase can be downloaded in PDB format accompanied by metadata (CSV format). An example depolymerase detail page is shown in Figure 1. The database information page features interactive statistical charts automatically generated from the current JSON file content. These include a donut chart showing the distribution of structural classes and subclasses (with the ability to explore subclasses by hovering over a given class segment), a bar chart illustrating K/KL type coverage, and pie charts displaying the distribution of phage morphotypes, life cycles, and phage genera.

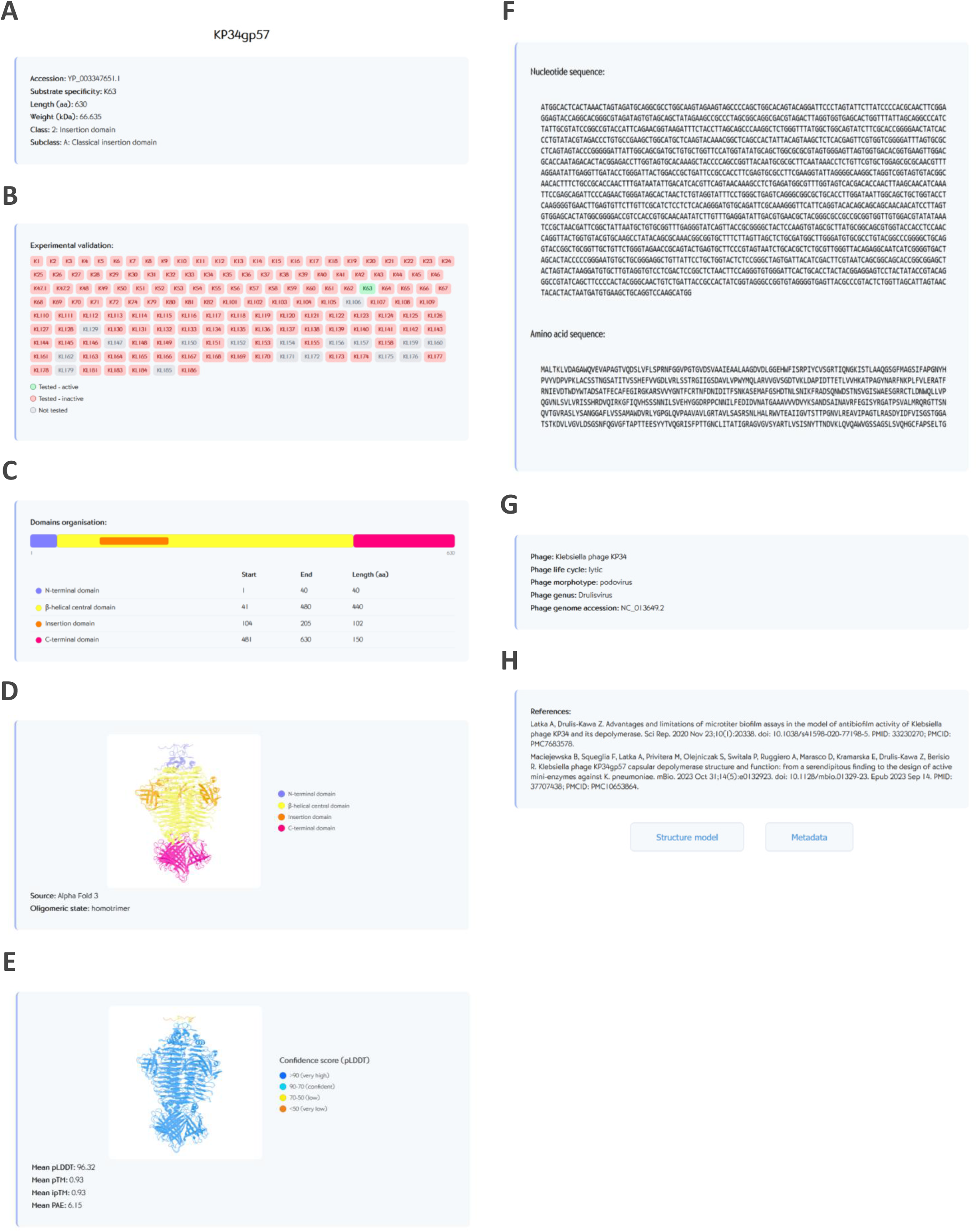

### Database content and statistics

The DepoCat database (v2.0) contains 131 experimentally validated Klebsiella phage depolymerases (Supplementary Table S1), covering activity against 75 distinct capsular types. Analysis of the distribution of entries by structural class revealed a predominance of Class 1 (classical depolymerase), comprising 80 proteins (61.1%) divided into six subclasses (1A–1F); Class 2 (insertion domain) accounts for 23 proteins (17.6%) across four subclasses (2A–2D); Class 3 (tail fiber domain) contains 17 proteins (13%) in two subclasses (3A–3B); Class 4 (α-helix-containing central domain) comprises 8 proteins (6.1%) in two subclasses (4A–4B); and Class 5 (colanidase-like depolymerase) contains 3 proteins (2.3%). The full distribution of structural classes and subclasses is presented in Figure 2A. Among the phage morphotypes – where the enzyme originates - podovirus (46 proteins, 35.1%) and jumbo myovirus (40 proteins, 30.5%) predominate. The vast majority of depolymerases are derived from lytic phages (111 proteins, 84.7%), while the remaining 20 proteins (15.3%) are of prophage origin. Among the phage genera, *Przondovirus* is the most represented (29 proteins, 22.1%), followed by *Maaswegvirus* (23 proteins, 17.6%) and *Drulisvirus* (13 proteins, 9.9%). The full distribution of phage morphotypes, life cycles, and phage genera is presented in Figure 2B.

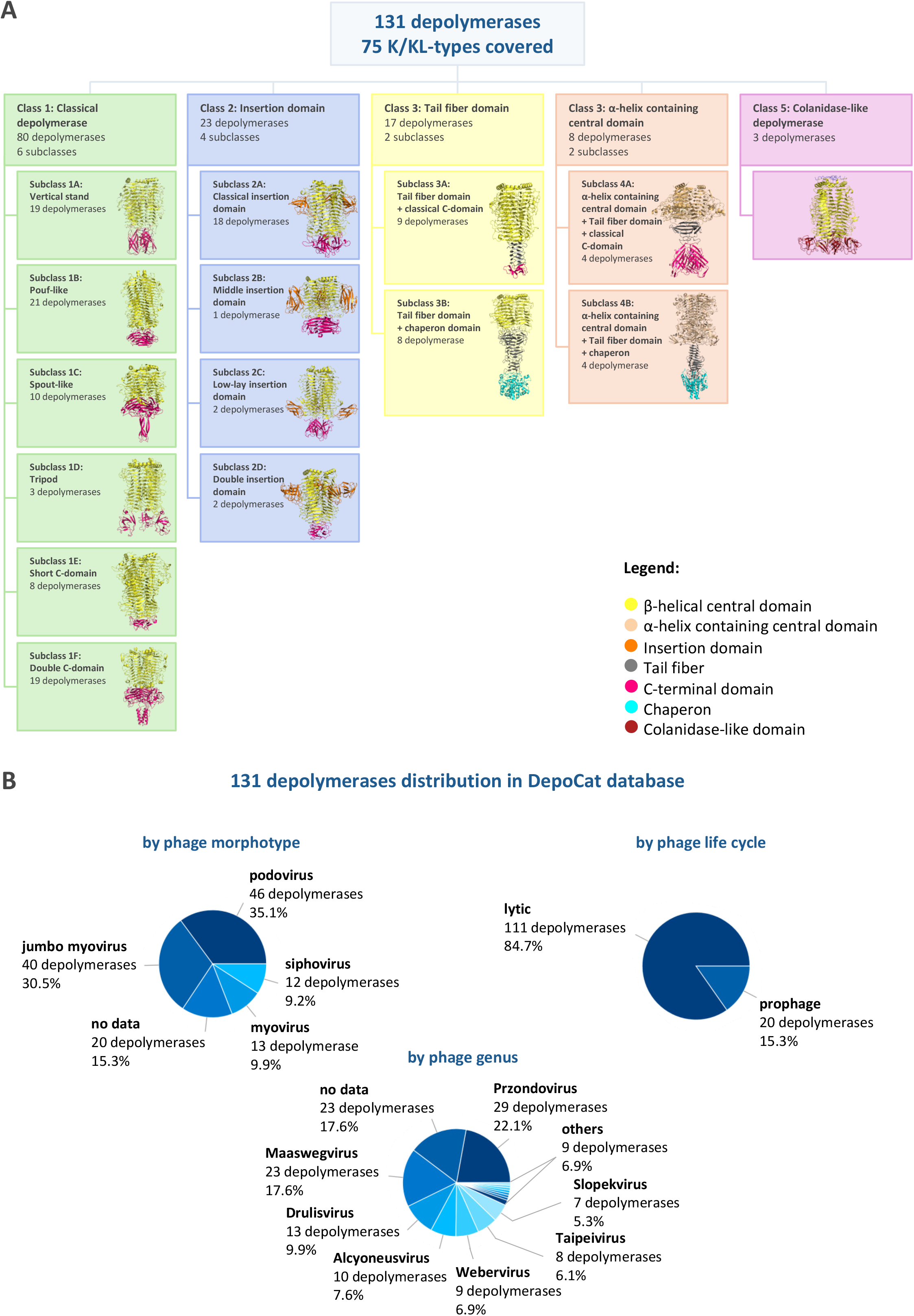

### Classifier tool: sequence search

The sequence search enables exploring the database using the amino acid sequence of a query protein, either pasted directly into a text field or uploaded from a FASTA-format file. BLASTP search results are presented in a sorted table containing, for each hit, values for sequence identity, query coverage, E-value, bit score, query and reference sequence lengths, alignment length, and the number of identical amino acid residues, together with depolymerase metadata including substrate specificity, structural class, and subclass. A graphical query coverage map displays results for all hits, colour-coded by percentage sequence identity: above 90% in red, 90–70% in orange, 70–50% in yellow, and below 50% in green. An alignment block showing the query and reference sequences alongside a consensus line is available for each hit. Supplementary Figure S1 shows example search results for a selected query protein.

### Classifier tool: structure search

The structure search allows users to upload a protein structural model in PDB or CIF format and compare it against all 131 reference structural models in the DepoCat database. Comparison results are presented in a table listing the TM-score for each match; this score ranges from 0 to 1, with 1 implicating an identical structural match and a score above 0.5 indicating that the structures generally share the same global topology. Scores below 0.17 correspond to randomly selected, unrelated proteins (35). The TM-score is reported in two variants: TM1 (normalised to the length of the query structure) and TM2 (normalised to the length of the reference structure). Additional results include RMSD (root mean square deviation, a metric quantifying the distance between atoms of superimposed molecules, expressed in Ångströms [Å]), alignment length, percentage sequence identity, and DepoCat metadata such as substrate specificity, structural class, and subclass. An interactive visualisation of the structural superposition is available for each match in a modal window, allowing full user interaction. Supplementary Figure S2 shows an example search result with the structural superposition.

### DepoCat beyond Klebsiella phage depolymerases

To illustrate the versatility of the DepoCat Classifier tool, we analysed three pairs of experimentally validated depolymerases with the same structural organisation but different substrate specificities and bacterial targets (Figure 3). The first pair, LKA1gp49 and phi297gp61, originates from two different phages infecting the same bacterial host (*Pseudomonas)* and acts on the same substrate as a lyase (LPS O5). These enzymes exhibit both detectable sequence similarity (76% coverage, 32.8% identity) and a nearly identical structural fold (TM-score = 0.95, RMSD = 1.901 Å). The second pair, LKA1gp49 and KP34gp57, comprises enzymes with distinct substrate specificities (LPS O5 lyase vs. CPS K63 hydrolase) and different origins (*Pseudomonas* phage vs. Klebsiella phage). No significant sequence similarity was found for this pair, although structural similarity was retained (TM-score = 0.69, RMSD = 5.473 Å). The third pair, KP34gp57 hydrolase and K1 lyase, originates from phages infecting the same bacterial host (*Klebsiella)*, yet the enzymes recognise distinct capsule types (CPS K63 vs. CPS K1) and possess different catalytic pocket locations. As in the previous case, no significant sequence similarity was found, while structural similarity was maintained (TM-score = 0.66, RMSD = 6.020 Å).

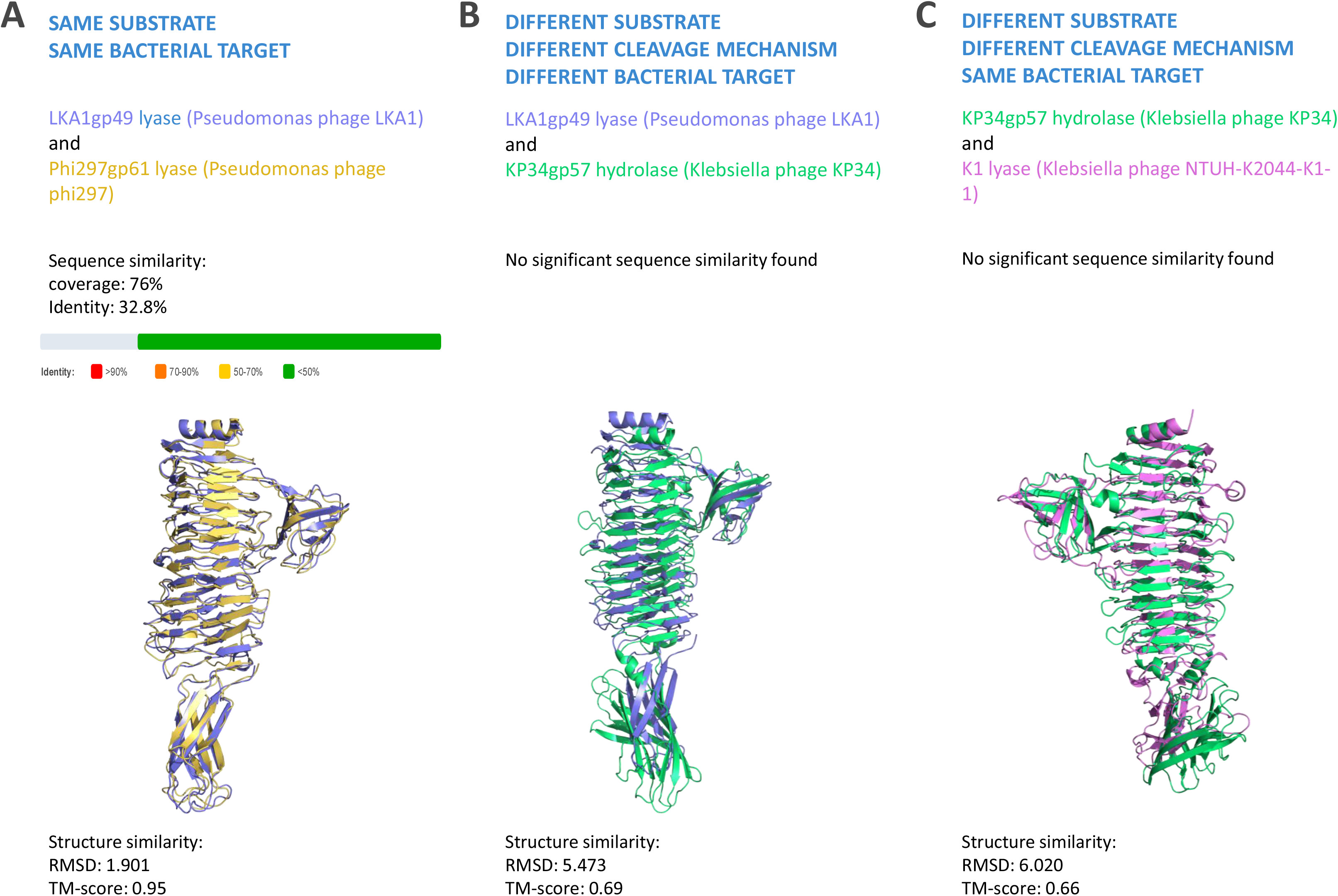

To determine whether this pattern holds across a broader, taxonomically diverse set of depolymerases, we further analysed 17 experimentally validated enzymes from *Acinetobacter, Escherichia, Pseudomonas, Salmonella and Shigella* phages (39–57)(Supplementary Table S2), comparing them at both sequence and structural levels against 131 reference Klebsiella phage depolymerases in the DepoCat database. TM1 values (normalised to the query protein length) for the best structural match ranged from 0.545 to 0.895 (median ≈ 0.70), with sequence identity remaining low across all these alignments (7.2-24.8%; Supplementary Figure S3, Supplementary Table S3). The highest values were obtained for ORF213_TSP4 (TM1 = 0.895; Dep1979, K2, Class 1B), DettilonTSP (TM1 = 0.868; P560dep, KL169, Class 1A), and Sf6_TSP (TM1 = 0.819; FKANgp220, KL148, Class 1A). The lowest were obtained for APK09gp48 (TM1 = 0.564; 434_33, K52, Class 3) and ORF211_TSP2 (TM1 = 0.622; 1723_59, K60, Class 1B). Structural classification was fully consistent among the top three hits (same class and subclass) for 7 of the 17 queries (HK620_TSP, K5 lyase KflA, LKA1gp49, ORF213_TSP4, phiAB6_TSP, phi297gp61 and Sf6_TSP), partially consistent (2 of 3 hits) for 8 queries (APK09gp48, APK14gp49, APK16gp47, Det7tsp, ORF210_TSP1, ORF211_TSP2, P22_TSP), and inconsistent for the remaining two queries (DettilonTSP and ORF212_TSP3), for which the top three hits represented different structural classes. Among these, ORF213_TSP4 provided the most unambiguous result overall, combining the highest TM1 value with high-quality superposition across all three reference hits. Among the remaining proteins with fully consistent classification, several exhibited good structural agreements within the central β-helical domain alongside weaker agreement in the C-terminal domain, as observed for LKA1gp49 (TM1 = 0.760-0.767), phi297gp61 (TM1 = 0.752-0.753) or Sf6_TSP (TM1 = 0.785–0.819). For LKA1gp49 and phi297gp61, all three top hits pointed to Class 2A “classical insertion domain”. For phiAB6_TSP (TM1 = 0.662–0.669), despite lower TM-score values and incomplete alignment in the direct superposition of the C-terminal domain, the overall domain topology – specifically the presence of a double C-terminal domain – corresponded to the subclass assigned to all three reference hits (Class 1F “double C-domain”). A distinct case was K5 lyase KflA, for which relatively lower TM-score values (0.594-0.690) were nonetheless accompanied by fully consistent classification of all three hits to the same class and subclass (Class 3B “tail fiber + chaperone”), matching the domain architecture of the query protein itself, which contains both a tail fiber domain and a chaperone domain in the C-terminal region.

The best sequence hits exhibited identities ranging from 20.4% to 76.2% and coverage of the query protein length ranging from 2.9% to 54.4% (Supplementary Figure S4, Supplementary Table S4). For three proteins – APK09gp48, APK16gp47, and P22_TSP – only a single statistically significant sequence hit was recorded across the entire reference set. The strongest sequence signals, in terms of both identity and coverage, were observed for ORF213_TSP4 (K7dep, Class 3B, 60.2% identity, 39.7% coverage), DettilonTSP (K27dep, Class 4B, 65.4% identity, 26.1% coverage), and ORF211_TSP2 (K27dep, Class 4B, 71% identity, 18.9% coverage). For the remaining 11 proteins with at least two hits, the best matches exhibited moderate identiity (22.3-52.9%) with low coverage (typically below 20%, except for Sf6_TSP, which reached 53.1% coverage for its best hit). In several cases – e.g. the third hit for phiAB6_TSP (76.2% identity, 2.9% coverage) – high identity was accompanied by extremely low coverage.

The class indicated by the best sequence hit agreed with that indicated by the best structural hit for 7 of the 17 proteins (9NA tailspike, HK620_TSP, K5 lyase KflA – partially, ORF210_TSP1 – partially, ORF212_TSP3, P22_TSP, Sf6_TSP), whereas discrepancies were observed for the others. Most notably for DettilonTSP and ORF211_TSP2, where the best sequence hit indicated Class 4B, while the best structural hit indicated Class 1A and Class 1B, respectively.

## Discussion

A key feature distinguishing DepoCat from many existing bioinformatics resources that gather protein data is its requirement for experimental validation of every entry. Large sequence repositories such as GenBank or UniProt contain numerous proteins annotated as “depolymerase” based on sequence homology or domain prediction, and the scale of this unvalidated functional diversity becomes apparent when searching for phage tail protein-related terms. As of August 25, 2026, queries combining “depolymerase” with “phage” in GenBank return nearly 17 946 records, with broader terms such as “tail spike” and “tail fiber” yielding 199 432 and 632 897 sequences, respectively. UniProt contains comparably lower but still substantial numbers, with 963 depolymerase-associated records and 5553 and 16 732 entries annotated as tail spike and tail fiber proteins, respectively. When restricted to Klebsiella phages, GenBank alone lists 15 255 sequences annotated as depolymerases, of which DepoCat (v2.0) includes 131 – exclusively those supported by experimental evidence of enzymatic activity and confirmed substrate specificity toward defined K/KL-types. UniProt similarly contains 381 records attributed to Klebsiella phage depolymerases, yet the majority of these annotations are derived computationally from sequence homology or domain architecture.

A similar gap exists among resources specifically dedicated to phage-derived proteins. The closest conceptual analogue to DepoCat is the PhaLP database (58,59), which compiles phage lysins, enzymes that degrade bacterial cell walls, together with sequence and structural metadata. While PhaLP serves as a comprehensive tool for lysin researchers, DepoCat plays a complementary role as a resource for depolymerases; a key distinguishing feature of DepoCat is its requirement for experimental validation of every entry. Several complementary resources should also be mentioned among the computational tools dedicated to phage depolymerases. DepoScope is a tool for the automated annotation and delineation of depolymerase domains using large language models, enabling identification of depolymerase candidates in genomic data at the amino acid sequence level (60). DposFinder extends these capabilities by simultaneously predicting the depolymerase and its corresponding host capsular type using a transformer architecture based on the ESM-2 model (61). PhageDPO represents another computational approach to identifying phage depolymerases, based on machine learning (62). However, the aforementioned tools operate on computational predictions, which lack experimental verification. DepoCat, presented in this study, plays a complementary role to aforementioned tools as an experimentally validated reference database, enabling the assessment and verification of computational predictions against confirmed data on activity and substrate specificity; thus, it can be further used for other databases improvement.

The Classifier tool enables searching of the database using amino acid sequences and structural models. High sequence similarity to validated DepoCat entries can help identify the substrate specificity of a query protein. Previous studies have shown that depolymerases exhibiting high sequence identity combined with high sequence coverage often display similar or identical substrate specificity. However, depolymerases with identical specificity can also exhibit varying degrees of sequence similarity (12). For this reason, the sequence search is complemented by a structure search. It enables comparison of a structural model query against the reference database even when sequence similarity is too low for effective BLASTP searching. TM-score values above 0.5 indicate shared global topology (35), serving as a basis for preliminary assignment to a structural class. A TM-score exceeding 0.9 suggests a similar or identical substrate specificity profile (12). However, a significant limitation of this method must be highlighted: the US-align program compares only the positions of Cα atoms in the peptide backbone, disregarding side-chain conformation, surface charge distribution, protein surface hydrophobicity, and active-site geometry entirely. These are precisely the features that decisively determine a depolymerases substrate specificity, as recognition of the polysaccharide substrate relies heavily on surface complementarity and electrostatic interactions between the enzyme and the substrate (8). Consequently, high similarity in peptide backbone does not categorically predict shared substrate specificity. Structural classification results should therefore be interpreted primarily as an assignment to a structural class; inferring substrate specificity from them is possible only to a limited extent and requires further analysis and experimental validation. These limitations stem from the current state of knowledge regarding sequence–structure–specificity relationships in this group of enzymes and define the directions for further development of the database and its tools.

Interpreting the results of the sequence analysis requires considering the relationship between sequence and structure. Above ∼40% identity in long alignments, a structural relationship is virtually certain, whereas below ∼20-25% identity - particularly in short alignments - does not constitute reliable evidence of a shared fold (63). This is particularly relevant to the present analysis, as DepoCat structural classes are defined solely on the basis of the architecture of the central and C-terminal domains (12). The N-terminal domain responsible for anchoring depolymerases to the phage baseplate/tail and for attaching other depolymerases within tail spike branching system (23), does not factor into class assignment. For most analysed proteins, the strongest sequence matches were confined to the N-terminal domain and were therefore uninformative for structural classification, irrespective of their statistical significance. This is best illustrated for DettilonTSP and ORF211_TSP2: the strongest sequence signal (matching K27dep/ORF96, Class 4B) covered only the first ∼150-200 residues of the protein, whereas the match spanning the central and C-terminal regions appeared only as the third hit, at lower sequence identity (P560dep, Class 1A).

Structural analysis makes it possible to determine which of these sequence signals genuinely reflect membership in the correct class. For DettilonTSP and ORF211_TSP2, it was the central and C-terminal domains – rather than the statistically stronger N-terminal alignment – that corresponded to the structural comparison results, confirming their classification into Classes 1A and 1B, respectively. For ORF213_TSP4, sequence-structure agreement in the central domain was supported by the most unambiguous result of the entire analysis (TM1 = 0.895, indicating near-complete structural similarity of the whole protein). Taken together, the results of both analyses indicate that the DepoCat Classifier tool applies correctly beyond its original scope of Klebsiella phage-derived depolymerases: for 7 of the 17 proteins, all three top structural hits indicated the same class and subclass, with TM1 values (0.545-0.895) ranging from moderate to high despite low sequence identity (7.2-24.8%) in every instance.

Notably, the absolute TM-score value was not always the sole indicator of classification confidence. K5 lyase KflA, despite relatively low values (0.594-0.690), yielded a fully consistent result, with all three hits pointing to the same class and subclass (3B: Tail fiber domain), matching the protein’s actual domain architecture. PhiAB6_TSP, despite imperfect agreement in the direct superposition of the C-terminal domain, exhibited an overall domain topology in this region (a double C-terminal domain) consistent with its assigned subclass (1F: Double C-domain). A particular illustrative case is ORF210_TSP1, for which structural comparison revealed partial similarity to Class 2 “Insertion domain” and Class 1 “Classical depolymerase” reference proteins alongside an atypical C-terminal domain topology not represented among current DepoCat entries, suggesting that this protein may constitute a structural variant corresponding to a novel subclass within Class 1.

The observed pattern of differential structural conservation across protein domains has broader evolutionary implications. Cases in which strong structural agreement is confined to the central β-helical domain while alignment in the C-terminal domain is weaker or incomplete – as observed for APK09gp48 or P22_TSP – may reflect the independent evolutionary trajectories of these two regions. This is consistent with the modular organisation of phage tail proteins, in which individual domains can evolve and disseminate independently, through horizontal gene transfer (HGT)(24). Under this model, the central domain, which forms the structural scaffold of the depolymerase, would represent a more conserved element, while the C-terminal domain – directly responsible for substrate recognition (10) – would be subject to more rapid changes driven by the selective pressure of capsular diversity.

More broadly, structural classification remains the primary and most reliable criterion for assigning a depolymerase to its correct architectural category. Sequence analysis can serve as a useful supplement only when high identity is accompanied by substantial coverage spanning the central and C-terminal domains. In other cases – particularly where alignments are restricted to the N-terminus, regardless of their apparent statistical significance – only structural comparison enables correct classification, as it detects the conserved fold even in the complete absence of sequence similarity in that region. Discrepancies in classification for some proteins may indicate that they represent structural variants not yet captured by the current DepoCat reference set – particularly given that the database currently comprises only 131 validated depolymerases from Klebsiella phages – pointing to a clear direction for future development: expanding the reference set to include validated depolymerases from a broader host range and identifying new structural subclasses. This complementarity between methods, combined with the primacy of structure as the decisive criterion, underlies the value of DepoCat as a universal tool for classifying phage depolymerases.

In summary, DepoCat is the first database dedicated to experimentally validated Klebsiella phage depolymerases, integrating experimental and computational data into a single, publicly accessible resource. It will serve both as a tool for the preliminary classification of newly described depolymerases and as a platform for systematic research into the structural diversity and evolution of this group of enzymes. The Classifier tool available via the web interface enables researchers to quickly access the database’s resources for their own research projects. Beyond its current scope, DepoCat is intended as a starting point and cornerstone for an integral and expanding database of experimentally validated phage depolymerases, irrespective of host species or targeted polysaccharide, supporting future research into depolymerase biology and application.

## Data availability

DepoCat data can be freely downloaded in various formats (depolymerase metadata in CSV, structural models in PDB) without restriction from http://depocat.uwr.edu.pl, provided that the origin of the data is cited when used in other work. No registration is required to access the data.

## Code availability

The source code of the DepoCat web application is not publicly available due to ongoing development; code snippets and analysis scripts may be made available to interested researchers upon reasonable request to the corresponding author.

## Acknowledgements

The DepoCat web application was developed with the assistance of Claude (Anthropic, claude.ai), an AI-based coding assistant.

## Author contributions statement

Conceptualization: SO, ZDK

Data Curation: SO

Formal Analysis: SO, AO, MP

Funding Acquisition: ZDK

Investigation: SO, AO, MP

Methodology: SO, AO

Project administration: ZDK

Resources: ZDK

Software: SO

Supervision: ZDK

Validation: SO

Visualization: SO, AO, MP

Writing – Original Draft: SO, ZDK

Writing – Review & Editing: all authors

